# Turquoise killifish naturally develop hallmarks of age-related macular degeneration with advancing age

**DOI:** 10.1101/2025.10.23.683644

**Authors:** Nicole C. L. Noel, Eva-Maria Breitenbach, Ryan B. MacDonald

**Affiliations:** Institute of Ophthalmology, University College London, London, UK

**Keywords:** age-related retinal disease, photoreceptors, retinal pigment epithelium, ageing, *Nothobranchius furzeri*

## Abstract

Ageing is a major risk factor for developing vision loss diseases such as age-related macular degeneration (AMD). Unfortunately, we do not have the ability to effectively prevent, slow, or stop onset and progression of AMD long term. These challenges with therapeutic development result from poor understanding of disease mechanism and pathogenesis due to a lack of animal models that manifest the hallmark features of disease. Here, we investigated the rapidly ageing turquoise killifish (*Nothobranchius furzeri*) retina for features of human ageing and AMD. We report that the ageing killifish retina expresses genes associated with human retinal disease in the photoreceptors and retinal pigment epithelium (RPE). Our characterisation of the retina identified that killifish spontaneously develop many hallmark features of AMD and human ageing, including photoreceptor deterioration, lipid deposits, outer retinal inflammation, and ceramide accumulation in the RPE with advancing age. Further, we identify a sex-specific difference in the severity of phenotypes. We propose that the turquoise killifish is a highly suitable model for investigating ageing and AMD-related disease mechanisms across the lifespan.

## Introduction

Age-associated ocular diseases such as age-related macular degeneration (AMD) are major causes of irreversible vision loss. Unfortunately, there are no treatments available to halt the progression of neurodegeneration in these conditions long term. Age-related conditions are challenging to study in patients, as they must be observed over decades, and are ideally screened early, to capture pathological changes associated with the onset and progression of disease. Further, tissue- or cellular-level resolution is challenging and not be feasible to acquire in patients as there is a scarcity of suitably preserved postmortem tissues to enable molecular investigation. Therefore, it is essential to identify model systems that recapitulate human disease to determine the pathological mechanisms underlying disease and to screen potential therapeutics.

The retina is the light-sensitive tissue found at the back of the eye. Degeneration of the light-detecting photoreceptor cells and retinal pigment epithelium (RPE), which are essential retinal support cells, result in vision loss in many retinal diseases (such as retinitis pigmentosa, macular degeneration, and early-onset severe retinal degeneration). Therefore, maintaining healthy photoreceptors and RPE throughout life is essential for maintaining optimal vision. AMD is further characterized by deposits at the back of the eye, retinal inflammation, and progressive degeneration of the cone photoreceptor cells and RPE in the central macula (Bergen et al., 2019; Curcio et al., 2005; Spaide et al., 2018), the specialised area for high-acuity vision (Figure 1). The hallmark deposits that increase risk of developing AMD are drusen and subretinal drusenoid deposits. Drusen are lipid-rich deposits between the RPE and underlying vessels that can also contain oxidised phospholipids, which indicate an oxidative stress environment and can trigger an immune response, potentially contributing to the inflammation observed in the AMD retina (Handa et al., 2017; Spaide et al., 2018). Subretinal drusenoid deposits are a lipid-containing (but not lipid-rich) granular deposit between the photoreceptor OSs and the RPE (Chen et al., 2021; Rudolf et al., 2008; Spaide et al., 2018). AMD is a complex condition, caused by a combination of environmental and genetic factors, but the biggest risk factor for developing AMD is ageing (Fritsche et al., 2013; Smith et al., 2001; Thornton et al., 2005). Unfortunately, we do not have complete understanding of why ageing increases likelihood of developing retinal disease; the cellular and molecular features that create an environment that is sensitized to degeneration require significant investigation. This poor understanding ultimately inhibits our ability to generate effective preventative therapeutics.

**Figure 1.**
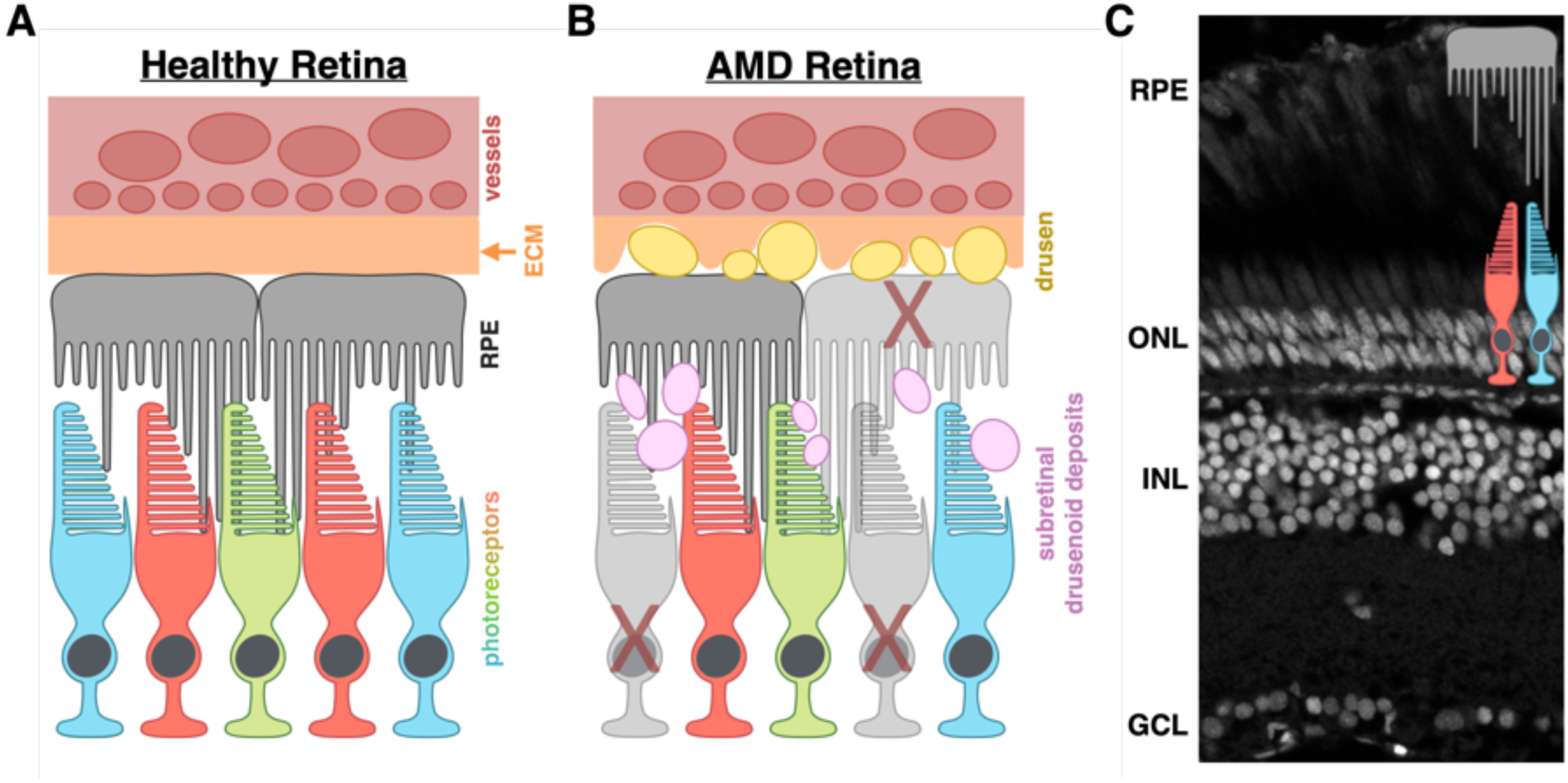
Age-related macular degeneration features. (A) Healthy and (B) AMD retinas. AMD retinas have the formation of drusen and subretinal drusenoid deposits, as well as degeneration of photoreceptors and the RPE. (C) Turquoise killifish retinal architecture as observed by DAPI staining on cryosectioned tissue, highlighting where the primary cell types of interest for these studies — the photoreceptors and RPE — are located. RPE = retinal pigment epithelium. ONL = outer nuclear layer; INL = inner nuclear layer; GCL = ganglion cell layer.

Further confounding the many challenges in studying AMD is a lack of accurate models. Nocturnal rodents have few cones and infrequently develop AMD hallmarks (i.e. deposits) (Fletcher et al., 2014). Zebrafish have cone-rich retinas that undergo age-related retinal changes (Allison et al., 2010; Martins et al., 2022); however, zebrafish are relatively long-lived (3-5 years), which creates challenges for study of age-related diseases. An emerging model is the turquoise killifish (*Nothobranchius furzeri*), a rapidly ageing vertebrate that naturally develops hallmark features of human ageing and age-related disease(Bergmans et al., 2024; de Bakker & Valenzano, 2023; Kim et al., 2016; Reichard et al., 2022; Vanhunsel et al., 2021). The short-lived GRZ strain of killifish have a lifespan of six to eight months, facilitating ageing studies within a significantly shorter timeframe than traditional vertebrate models (Kim et al., 2016). Killifish have been shown to have vision loss, retinal inflammation, and retinal thinning with advancing age (Bergmans et al., 2024; Vanhunsel et al., 2021). Transcriptomics have been used to characterise the retinal cell populations and reported that killifish have age-related transcriptional dysregulation, whereby cells turned on or increased expression of genes not typically enriched for that cell population (Bergmans et al., 2024). However, detailed histological characterisation of the age-related pathologies within the killifish retina have not been explored. Additionally, only female killifish retinas have previously been characterised – studies have not assessed whether male and female killifish retinas are differentially impacted by age. As killifish are a sexually dimorphic species, they provide the opportunity to investigate sex-specific neurodegeneration phenotypes during ageing. It is therefore unclear whether killifish naturally develop pathologies similar to age-related retinal diseases and whether there are differences in ageing phenotypes between the sexes.

Here, we investigated the turquoise killifish retina for features of human retinal ageing and AMD, observing that there were shared presentations. Photoreceptors have trapping of opsin protein within the cell body and synapse, indicative of photoreceptor degeneration. Further, we determined that turquoise killifish develop hallmark features of AMD such as lipid-rich deposits, retinal inflammation, and ceramide accumulation in the RPE. We also observed that male killifish have more severe retinal phenotypes than females. In sum, we have found specific AMD-like pathologies in the turqoise killifish and propose it is a suitable model to study age-related retinal disease.

## Methods

### Animal Care

GRZ strain turquoise killifish (*Nothobranchius furzeri*) were housed at University College London under a 14:10 light:dark cycle on PPLs 70/7391 and PP2133797. Animals were hatched once they reached the golden eye stage (uniformly gold coloured irises) by placing the embryos in an aerated 1:1 mixture of 4°C hatching solution (1g/L humic acid in Ringers solution): room temperature system water. Adult fish (4 weeks post hatching) were fed dry food and bloodworm. For all experiments, “young” fish were considered 6-week-old young adults, and “old” fish were considered 24-week-old adults at the end of their lifespan.

### Tissue Preparation

Killifish eyes were enucleated and fixed in 4% paraformaldehyde + 5% sucrose overnight at 4°C. Fixed eyes were washed with PBS to remove any residual fixative, then cryoprotected in 30% sucrose overnight at 4°C. Cryoprotected eyes were embedded in 3% agarose then allowed to equilibrate in 30% sucrose until the agarose block sank. Agarose-embedded eyes were then transferred into Tissue Freezing Medium (Merck Life Sciences, Cat. No. SHH0024) and frozen solid. Cryoblocks were stored at -80°C until use. Blocks were sectioned on a Leica CM1950 cryostat at a thickness of 12-15µm. Tissue was allow to dry for minimum for 20 minutes at room temperature before being stored at -80°C until use. Slides with cryosectioned tissue were warmed to room temperature for 1 hour and rehydrated with PBS prior to histological staining.

### Oil Red O

After rehydration, slides were fixed in 4% paraformaldehyde for 20 minutes, rinsed with distilled water, then rinsed with 60% isopropanol. Slides were stained with oil red O working solution (0.3% in 60% isopropanol) for 15 minutes, then rinsed with 60% isopropanol. Nuclei were stained with hemotoxylin (5 dips) and rinsed with distilled water. Excess liquid was tapped off, then Fluoromount G Mounting Medium (Thermo Fisher, Cat. No. 00-4958-02) applied and a coverslip added. Slides were imaged on a Leica THUNDER imager.

### Immunofluorescence

Slides were blocked for 60 minutes in 10% normal goat serum (NGS) in PBS + 0.1% Tween + 1% DMSO (PBSTw+DMSO). Primary antibodies were diluted in 2% NGS in PBSTw+DMSO, then applied to the slides. Slides were incubated in antibody overnight at 4°C in a humidity chamber. Excess antibody was removed with three washes of PBS. Secondary antibodies were diluted in 2% NGS in PBSTw+DMSO, and relevant other stains added (DAPI, LipidTox), then applied to the slides, and incubated in a humidity chamber overnight at 4°C. Excess secondary and fluorescent stains were removed by washing three times with PBS. FluorSave Reagent mounting media (Merck Millipore, Cat. No. 345789) was then added and the slides coverslipped for imaging on a Zeiss LSM 900.

### *in situ* Hybridisation Chain Reaction

The *in situ* hybridisation chain reaction (HCR) methods were modified from previously published protocols (Choi et al., 2018). Slides were permeabilised with 30 µg/mL proteinase K for 10 min at room temperature followed by a PBS rinse and 20 minute postfix in 4% PFA. The slides were then pre-hybridised in probe hybridisation buffer (Molecular Instruments, www.molecularinstruments.com) at 37°C for 30 minutes and probes mixed to a final concentration of 20 pmol/mL in probe hybridisation buffer. Probe mix was added to slides and incubated at 37°C for 3 days in a humidity chamber. Excess probes were removed by washing with 37°C wash buffer (Molecular Instruments) 3 x 10 minutes, followed by 2 x 5 minute washes with SSCT at room temperature. Slides were pre-amplified with amplification buffer (Molecular Instruments) at room temperature for 30 minutes. Hairpins were separately heated to 95°C for 90 seconds and snap cooled in the dark for 30 minutes. Cooled hairpins were mixed in amplification buffer and applied to the slides then incubated in the dark at room temperature for three days in a humidity chamber. Slides were washed with SSCT 4 x 15 minutes, stained with DAPI, mounted using FluorSave Reagent (Merck Millipore, Cat. No. 345789), and imaged with a Zeiss LSM 900.

### Histological measurements

Photoreceptor measurements: The length of the inner and outer segment regions (top of the outer nuclear layer to the very tips of the outer segments) was measured on central retina sections. Inner segments were included in this measurement because the inner segments do not always form a clear delineation across the retina, especially in old individuals, whereas the nuclear layer stays relatively uniform. For outer segment width, photoreceptors were labelled with PRPH2 antibody. Outer segment width was measured for 10 rod outer segments per individual.

### Statistics

Statistical analysis was performed using Prism version 10.3.1 (GraphPad Software Inc., San Diego, CA, USA).

## Results

### Turquoise killifish have conserved expression of disease-associated retinal genes

Genes associated with inherited retinal diseases typically encode essential components for photoreceptor or RPE structure and function; when these are perturbed, cellular degeneration occurs (Bachmann-Gagescu & Neuhauss, 2019; Bujakowska et al., 2017; Karali & Banfi, 2015; Yang et al., 2021). We previously showed that the killifish retina consists of conserved cell types and that there is transcriptome dysregulation in specific cell types across ageing (Bergmans et al., 2024). However, the expression and localisation of genes that cause retinal dystrophy when mutated have not been well assessed in the killifish, and so we first assessed whether disease-associated genes are expressed in the outer retina. Few photoreceptor or RPE-related genes have been characterised in the killifish retina, other than genes related to cell identity (Bergmans et al., 2024). We utilised *in situ* hybridisation chain reaction (HCR) to determine the cell specificity of several retinal disease-associated genes and assess whether there was a change in expression pattern of these genes in the aged killifish retina, suggesting cell stress and/or dysfunction. We found that young, 6-week-old adult killifish retinas expressed *rpe65a*, *rp1l1*, *prph2b*, *elovl4b*, and *eys* and their expression localised to the expected cell types; the RPE expressed *rpe65a* while photoreceptors expressed all other genes (Figure 2). We observed that 24-week-old retinas similarly expressed these genes in the expected cell types. There may be some age-related changes in expression patterns for *rp1l1*, *prph2b*, and *rpe65a* in old fish, but overall, there was no observed change in cell types expressing disease-associated genes between young and old fish. Thus, killifish express retinal disease-related genes in the expected cell types based on other model systems and human expression data, highlighting conserved outer retinal features that allow them to be used as a potential model for disease.

**Figure 2.**
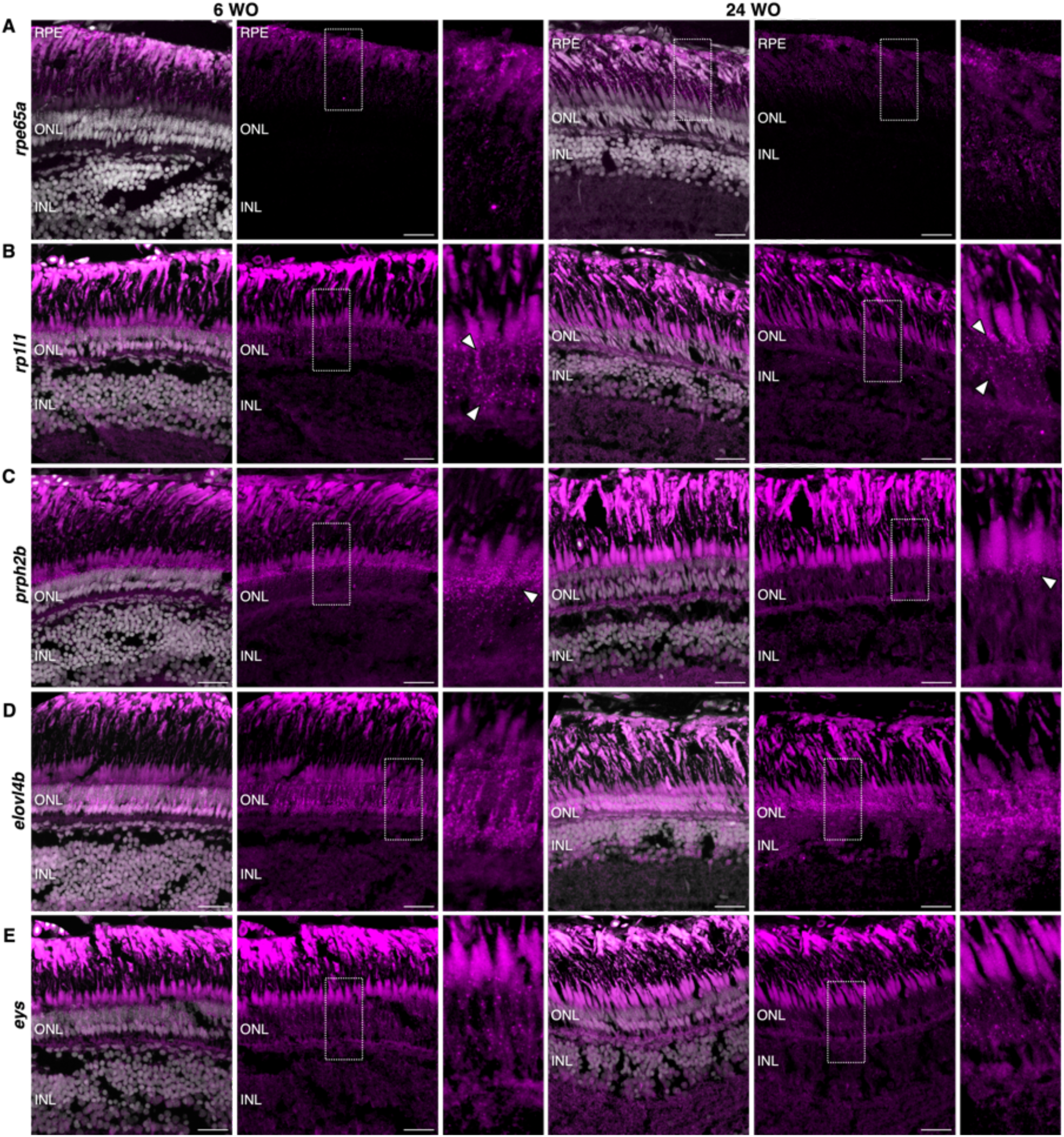
Outer retinal genes associated with inherited retinal degenerations are expressed in predicted cell types in young and old turquoise killifish. (A) rpe65a expression in young and old killifish is within the RPE. (B) rp1l1 expression is observed in photoreceptors (arrowheads) in young and old killifish. (C) prph2b expression is predominantly in cone photoreceptors (arrowheads). (D) Expression of elovl4b is largely restricted to rod photoreceptors. (E) eys expression is in photoreceptor cells in young and old killifish retina. Nuclei labelled with DAPI. Dotted boxes show magnified regions. INL = inner nuclear layer. ONL = outer nuclear layer. RPE = retinal pigment epithelium. INL = inner nuclear layer. ONL = outer nuclear layer. WO = weeks old. Scale bar = 25µm.

### Killifish photoreceptors have signs of degeneration with age

Next, we assessed whether killifish display features of human retinal ageing. Photoreceptor outer segments (OSs), the light-detecting part of the photoreceptor, are known to shorten with age in humans (Berlin et al., 2023; Bringmann et al., 2022; Cunea & Jeffery, 2007; Johnson et al., 2003). The OS sits above the inner segment (IS), a specialised compartment that sits below the OS and contains densely packed mitochondria. To first determine if killifish photoreceptors display the same features of age-related deterioration, we conducted histological analysis and measured the length of the photoreceptor IS/OS region within the central retina of male and female killifish at 6 and 24 weeks old (Figure 3A,B). We observed that 24-week-old male killifish had significantly shorter photoreceptor IS/OS regions than 6-week-old male and female fish (Figure 3C). Of note, while 24-week-old female killifish photoreceptors were not significantly shorter than 6-week-old killifish photoreceptors, they were trending towards being shorter, and the 24-week-old male killifish photoreceptors were not significantly different from the 24-week-old females. Since length of the IS/OS region could be artificially changing due to widening of the OSs rather than loss of photoreceptor material, OS width was also measured. There was no statistically significant difference in OS width between any of the groups (Figure 3D). Taken together, this suggests that male killifish have a loss of OS material with age, leading to shortening, rather than changes to gross OS morphology.

**Figure 3.**
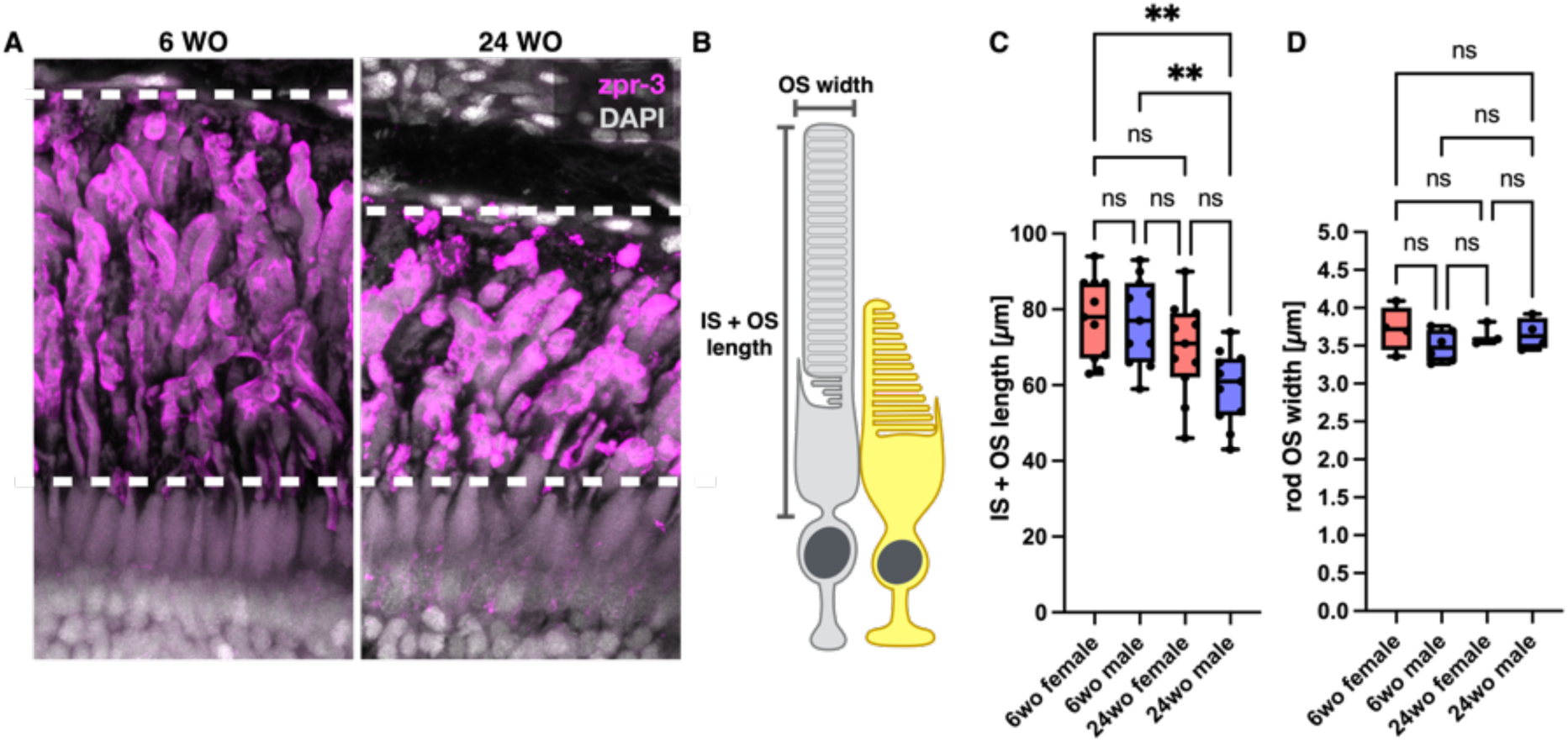
Age-related shortening of the photoreceptors in male killifish. (A) Fluorescent micrographs of young and old killifish retinas with photoreceptor outer segments labelled with zpr-3 showing shortening of the photoreceptor outer segments. (B) Cartoon of photoreceptor cells, rods (grey, left) and cones (yellow, left). For photoreceptor measurements, the entire inner and outer segment region was measured; for width measurements, width of rod outer segments were measured. (C) IS + OS measurements for 6- and 24-week-old female and male killifish. 24-week-old male killifish had significantly shorter IS + OS length compared to 6-week-old fish. n=10 per group. (D) Rod OS width was not different between the groups. n=6 for 6wo males; n=4 for 6wo females and 24wo males; n=3 for 24wo females. wo = weeks old; ns = not significant. **p<0.01, one-way ANOVA.

To assess changes in protein localisation, we labelled photoreceptor OSs to assess changes in protein localisation between young and old killifish. Zpr-3 and PRPH2 antibodies both label photoreceptor OSs; zpr-3 labels opsin proteins in rods and double cones while Prph2 is a membrane protein essential for establishment of OS disc morphology. In 24-week-old killifish, photoreceptors were observed with zpr-3 and PRPH2 labelling in the ISs, cell bodies, and synaptic terminal (Figure 4). Of note, these cells did not have an identifiable OS, indicating that the observed photoreceptor OS protein trapping was due to OS degeneration. These results demonstrate that aged killifish have photoreceptor OS deterioration, and taken together with the conserved expression patterns for photoreceptor OS genes (Figure 2), killifish are an eminently suitable model for age-related photoreceptor pathologies.

**Figure 4.**
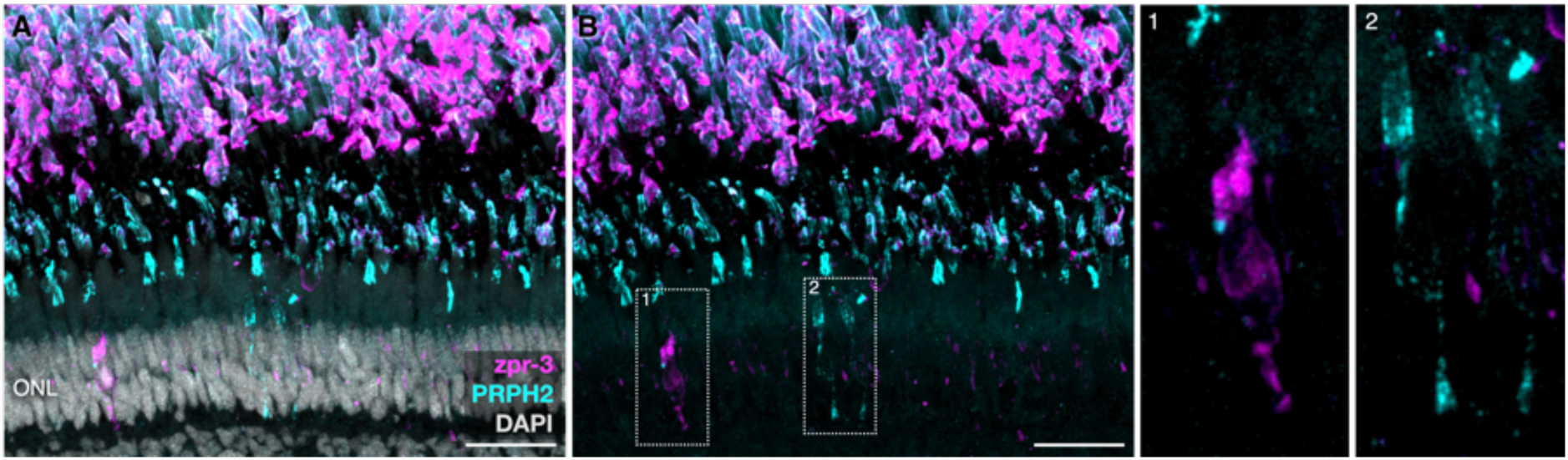
Old killifish have outer segment degeneration and trapping of outer segment proteins in the photoreceptor inner segment, cell body, and synaptic terminal. Photoreceptor outer segments labelled with zpr-3 and PRPH2, with (A) and without (B) DAPI on 24-week-old killifish retinal sections. Inset box 1 shows a cell with opsin protein trapped within the inner segment, cell body, and pedicle; inset box 2 shows cells with peripherin-2 (PRPH2) tapped in the inner segment, cell body, and pedicle. ONL = outer nuclear layer. OS = outer segments. Scale bars = 25µm.

### Aged killifish retinas have markers of RPE stress, including deposit formation and ceramide accumulation

A key hallmark feature of AMD is deposits – subretinal drusenoid deposits and drusen – which increase risk of developing AMD. To assess whether killifish naturally develop deposits as they age, retinas were stained for neutral lipids using LipidTox and oil red O (Figure 4). Deposits were mainly found in the central retina, with few deposits observed in the peripheral retina (Figure S1). Deposits were typically between the photoreceptor OSs and RPE but could also be observed sub-RPE (Figure 5A-C, S1). As there may be differences in incidence and severity of AMD between the sexes in humans (Owen et al., 2012; Rudnicka et al., 2012; Sasaki et al., 2018), we investigated whether there were sex differences in deposit accumulation in killifish. Counting the number of deposits within the central retina for male and female killifish revealed that male killifish had significantly more deposits than females (Figure 5D) – the average deposit number for males was >4-fold higher. In male retinas with large clusters of deposits, lipid deposits could be observed amongst the photoreceptor ISs, within the outer nuclear layer, and occasionally the inner nuclear layer (Figure 5C, S2). Deposits could be seen as early as 18 weeks of age in male killifish (Figure S3). Immunofluorescence with an antibody against oxidised phospholipids revealed that the deposits were positive for oxidised phospholipids (Figure 5A-B); in females, deposits were often positive for both neutral lipids and oxidised phospholipids, but in males, greater deposit diversity was observed with some deposits staining for only neutral lipids. Interestingly, what appeared to be oxidised phospholipid-rich OS fragments could be observed, which may be indicative of OS breakdown (Figure 5B). Opsin proteins labelled with zpr-3 antibody were observed within or closely surrounding lipid deposits (Figure 6). This could reveal how the deposits form or provide insight into how the deposits interact with OS proteins.

**Figure 5.**
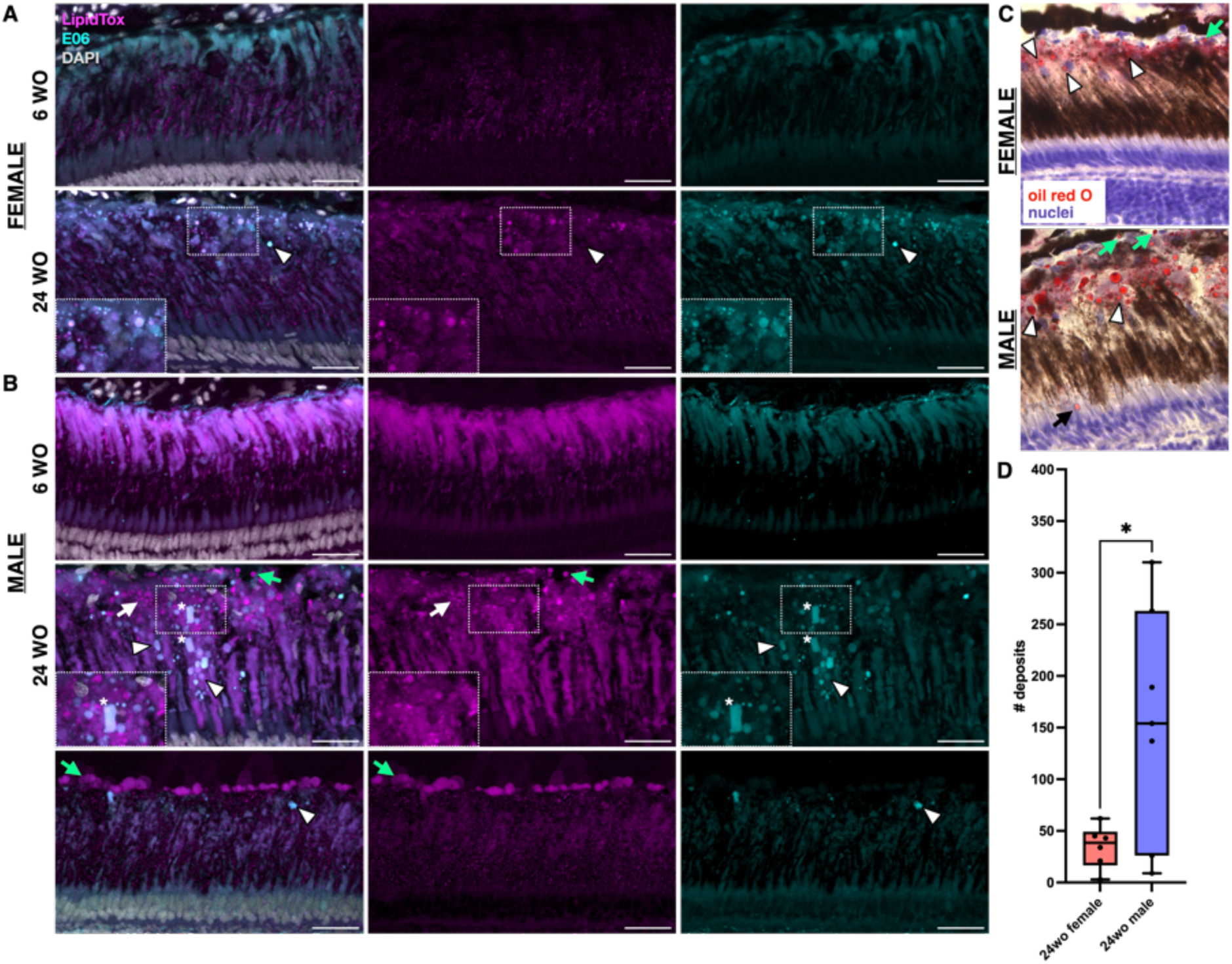
Aged killifish develop deposits on the apical and basal side of the RPE. (A-B) Confocal images of young (6-week-old) and old (24-week-old) killifish retinal sections with neutral lipids labelled with LipidTox and oxidised phospholipids labelled with E06 antibody. (A) Young (top) and old (bottom) female killifish retinas. Young retinas do not have visible deposits, while old retinas have clear deposits labelled (insets), including deposits highly enriched for oxidised phospholipids (arrowhead). (B) Young (top) and old (middle, bottom) male killifish retinas. 24-week-old males had deposits that were enriched for neutral lipids and oxidised phospholipids (insets, middle panels). Deposits were observed on both the apical (white arrows) and basal (green arrows) side of the RPE, including retinas where deposits were primarily basal (bottom panels). Deposits were enriched for oxidised phospholipids (arrowheads). There were oxidised phospholipid-rich outer segment-like fragments (asterisks). (C) Oil red O and hematoxylin staining on 24-week-old female (top) and male (bottom) killifish retinas, similarly showing subretinal lipid deposits (arrowheads) and lipid deposits on the basal side of the RPE (green arrows). Male retina had lipid deposits in the photoreceptor outer segment layer (black arrow). (D) Quantification of deposit number in 24 weeks old female and male retinas across a 225µm area. Males had significantly more deposits (155.4 ± 111.9) than females (34.7 ± 20.6). wo = weeks old. n=6 for females; n=7 for males. *p<0.05, Welch’s t test. OS = outer segments. Scale bars = 25µm.

**Figure 6.**
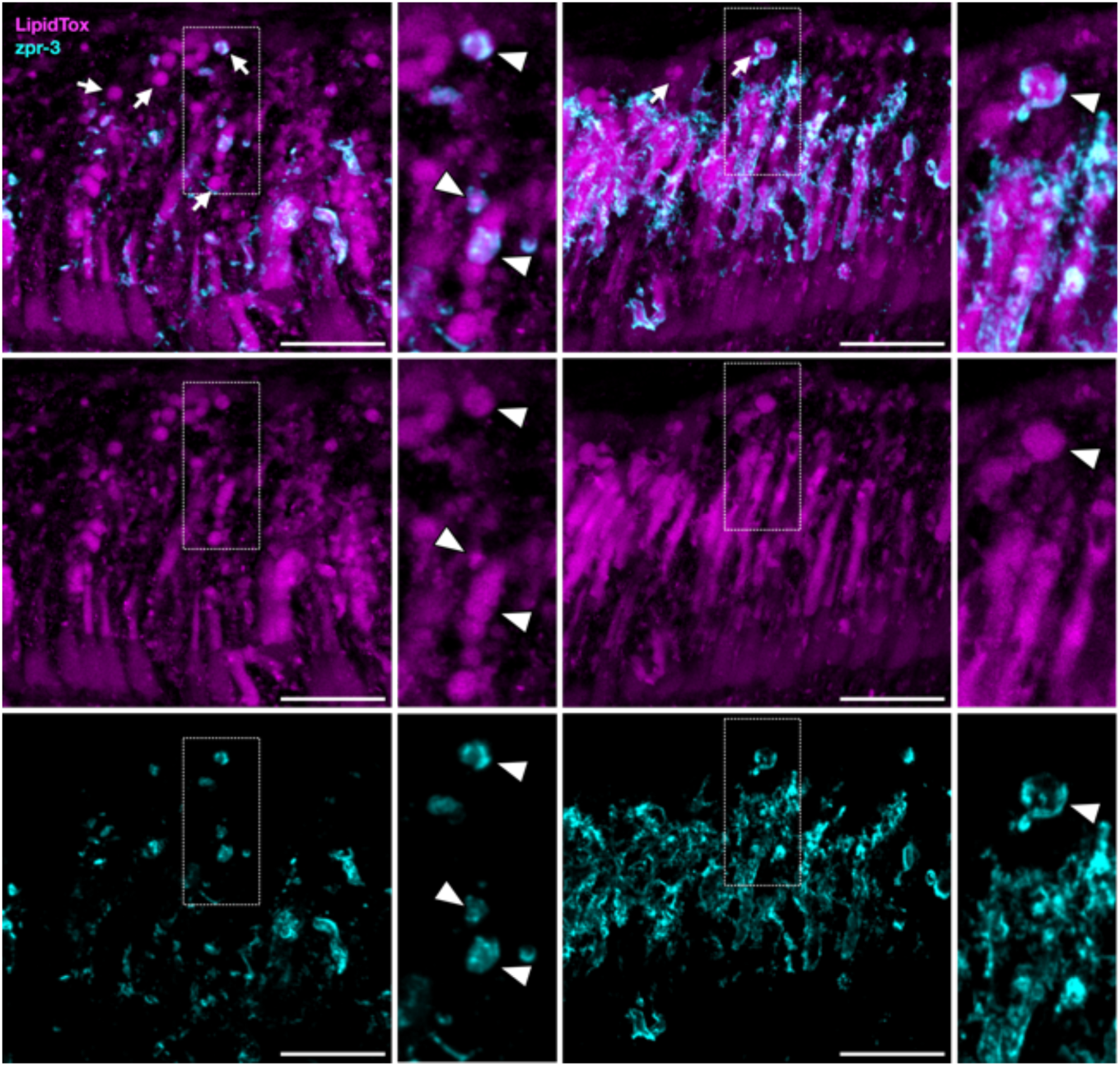
Lipid deposits can contain outer segment proteins. 24-week-old killifish retinas with neutral lipids labelled (LipidTox, magenta) and photoreceptor opsins (zpr-3, cyan) labelled. Lipid deposits contained opsin proteins (insets, arrowheads). Scale bars = 25µm.

Ceramide accumulation in the RPE indicates stress and dysfunction and can even trigger RPE cell death (Barak et al., 2001; Kaur et al., 2018). We assessed the RPE for additional signs of dysfunction and degeneration by staining for ceramide. We assessed the RPE for ceramide accumulation using immunofluorescence. In the 6-week-old killifish, ceramide was restricted to the neural retina, specifically associating with Müller glia (Figure 7; Figure S4). In contrast, in 24-week-old retinas ceramide was again observed within the neural retina but also accumulated in the RPE. Ceramide accumulation was not uniform between cells, and neighbouring RPE cells could show different amounts of labelling.

**Figure 7.**
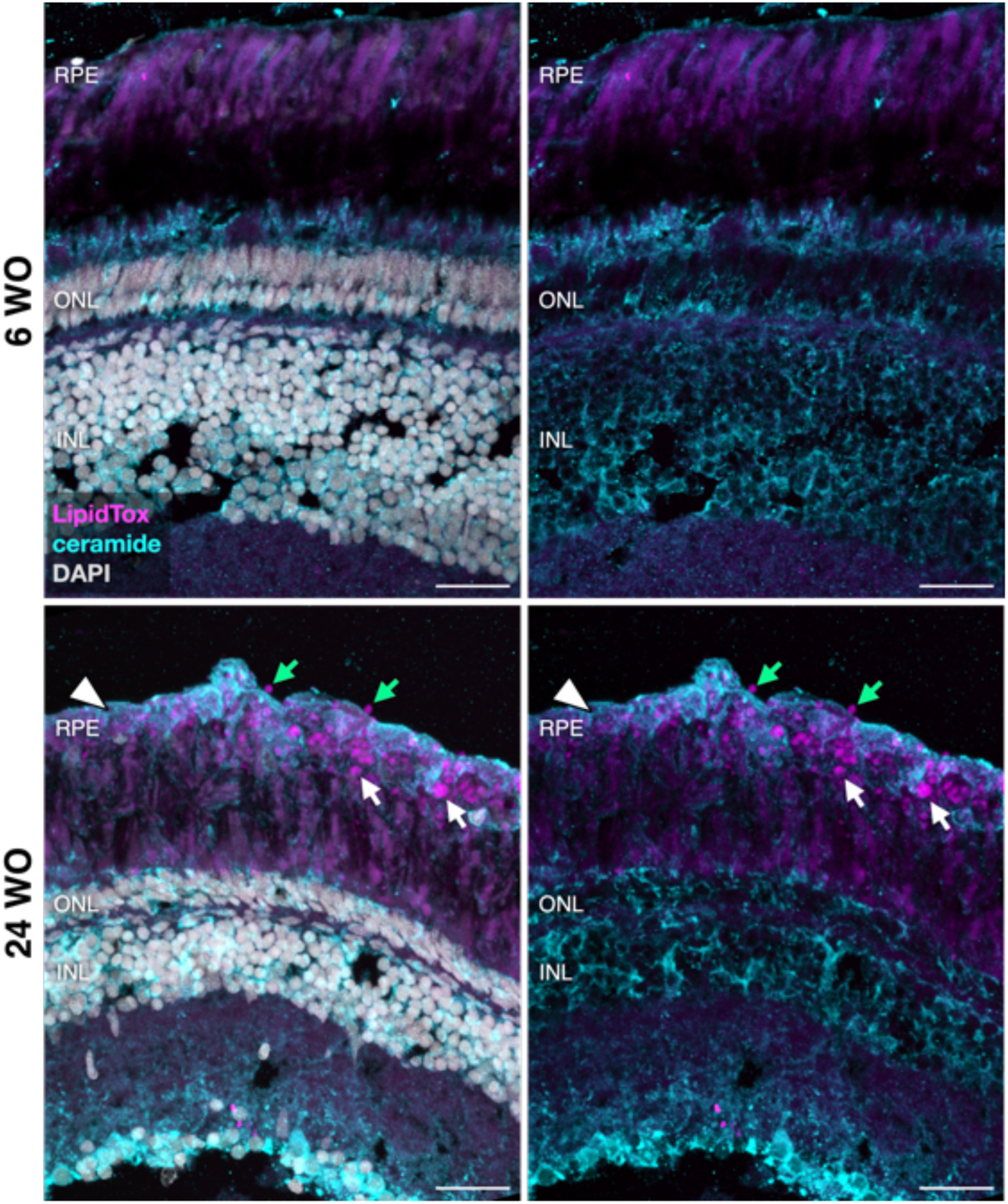
Ceramide accumulates in the retinal pigment epithelium of old killifish. Young (6-week-old) and old (24-week-old) killifish retinas labelled with ceramide using an antibody and neutral lipids were labelled via LipidTox with DAPI (left) and without (right). Old killifish have accumulation of ceramide (cyan) within the RPE (arrowhead), while young killifish have ceramide restricted to the neural retina. Lipid deposits (magenta) are observed between the photoreceptors and RPE (white arrows) as well as behind the RPE (green arrows). INL = inner nuclear layer. ONL = outer nuclear layer. RPE = retinal pigment epithelium. Scale bar = 25µm.

These results demonstrate that killifish develop lipid-rich deposits on the apical and basal side of the RPE. There are sex-specific differences in the timing and severity of the phenotypes, with male killifish developing deposits earlier. Additionally, aged killifish RPE has ceramide accumulation, indicative of stress and dysfunction.

### Killifish have macrophages in the subretinal space in areas with deposits

Killifish retinas have been reported to have increased macrophages with age (Bergmans et al., 2024; Vanhunsel et al., 2021) – however, whether there is translocation of macrophages in the subretinal space has not been investigated. As there are oxidised phospholipid-rich deposits between the OSs and RPE and oxidised phospholipids can trigger an immune response, we hypothesised that there would be an accumulation of macrophages in the subretinal space in old killifish. We performed immunofluorescence using an L-plastin antibody to label macrophages. 6-week-old male and female killifish had few macrophages in the OS region, and when they did have macrophages, the cells were in close proximity to the RPE and had a spherical cell body morphology with fine processes (Figure 8A). Conversely, 24-week-old male and female killifish had macrophages throughout the OS region, including macrophages that appeared to be wrapping around photoreceptor outer segments (Figure 8B). These macrophages had thick processes, did not have spherical cell bodies, and appeared motile. We counted the L-plastin+ macrophages the central retina in young and old male and female killifish as well as in the comparatively younger peripheral retina in old killifish, observing that old male killifish had significantly more macrophages in the subretinal area than young males and females (Figure 8C). Similarly, the central retinas of old males had significantly more macrophages than the comparatively younger peripheral retina. Old females had significantly more macrophages in their outer retina than young male central and old male peripheral retinas, but there was no significant difference between young female central or old female peripheral and old female central retina. In conclusion, aged killifish have translocation of macrophages to the outer retina, which coincides with AMD phenotypes, and male fish display more prominent macrophage phenotypes.

**Figure 8.**
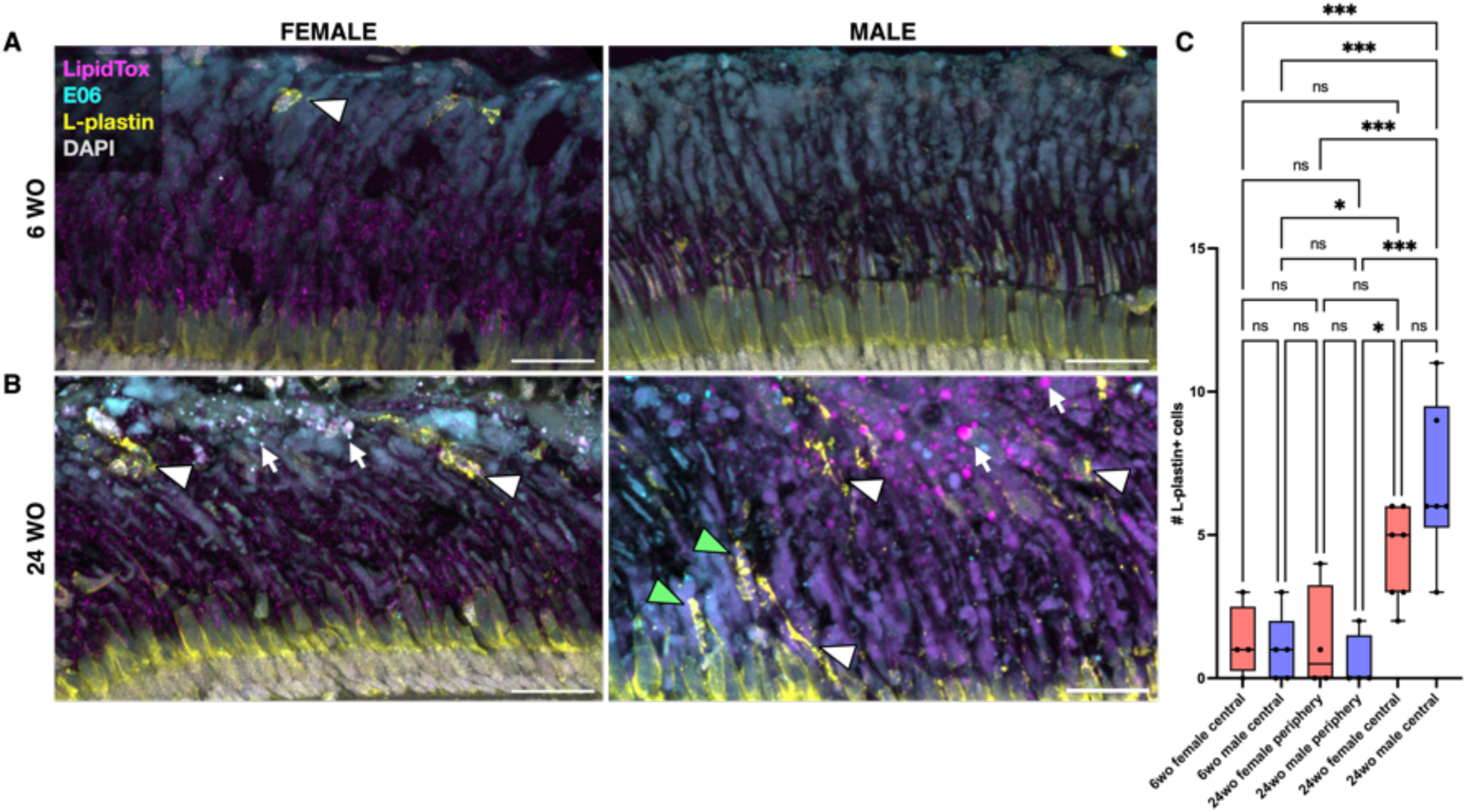
Aged killifish have increased macrophages in the subretinal space. Confocal images of young (A) and old (B) female and male killifish retinas labelled with LipidTox for neutral lipids (magenta), E06 for oxidised phospholipids (cyan), and L-plastin antibody (yellow). (A) Macrophages in the subretinal space of young fish sat towards the tips of the outer segments in close proximity to the RPE, with rounded somas and fine processes. (B) Macrophages in old retinas had expanded morphologyes and were observed throughout the subretinal space (arrowheads), with some appear to be wrapped around outer segments (green arrowheads). Deposits are visible in old retinas (arrows). (C) Quantification of the number of L-plastin positive cells within a 225µm area. *p<0.05; **p<0.01; ***p<0.001, one-way ANOVA. n=4 for 6wo female; n=5 for 6wo male and 24wo female; n=6 for 24wo male. ns = not significant. wo = weeks old. Scale bar = 25µm.

## Discussion

Ageing is the most significant risk factor for developing age-related retinal disease. Turquoise killifish, an emerging rapidly ageing model, allows for study of age-related pathologies within a short time frame. We investigated the histological presentations of the killifish retina between young and old male and female animals and found that killifish naturally develop hallmarks of age-related retinal disease, including AMD. Further, we show sex-specific differences in killifish retina ageing, with male killifish having earlier onset and more severe retinal phenotypes.

### Killifish display hallmarks of human ageing and age-related macular degeneration

There are many parallels between AMD features and the observed killifish retinal phenotypes reported here. The spontaneously occurring pathologies in the killifish retina that are similar to AMD may manifest in part due to photoreceptor composition; as a diurnal teleost fish, killifish have a cone-rich retina. This means that the killifish retina is more similar to the central macula of humans than the retinas of other species, such as nocturnal rodents, which have few cones. Killifish naturally develop deposits, a hallmark feature of AMD, on both the apical and basal side of the RPE. In humans, drusen typically form in the comparatively cone-rich central retina, while subretinal drusenoid deposits develop in more rod-rich areas (Bergen et al., 2019; Chen et al., 2021; Monge et al., 2022; Rudolf et al., 2008; Spaide et al., 2018); killifish may have the capacity to develop deposits on both the apical and basal sides of the RPE because they an abundance of rods and cones that contributes to this, unlike other species with rod-dominant retinas. Subretinal lipid deposits in killifish have features of both drusen and subretinal drusenoid deposits, as they are be located between the photoreceptors and RPE – where subretinal drusenoid deposits are found – but are lipid-rich and oxidised phospholipid-containing, which are features of drusen (Handa et al., 2017; Spaide et al., 2018). Further, the killifish retina has accumulation of oxidative stress with age (Bergmans et al., 2024; Vanhunsel et al., 2021), which is similarly observed in humans (Handa et al., 2017). Additionally, killifish have retinal inflammation with translocation of immune cells to the outer retina, also observed in AMD.

We do not understand why deposits form during ageing and AMD, nor how they develop. This is due to there being few opportunities to study deposit formation *in vivo*, as few animal models develop deposits, and those that do are typically mutant lines (Ban et al., 2018; Carr et al., 2024; Cheng et al., 2020; Imamura et al., 2006; Noel et al., 2020). It is possible that OS deterioration and/or RPE dysfunction may result in ineffective uptake of OS fragments and accumulation in the subretinal space, leading to deposits. Alternatively, RPE degeneration may be leading to RPE contents, such as opsin-containing phagosomes, being expelled into the subretinal space which subsequently breaks down. Killifish can facilitate the exploration of how the phagocytic capacity and cellular stability of the RPE changes with ageing to address these questions.

It is not known why photoreceptor OSs shorten with age. The killifish can be used to assess whether old photoreceptors have perturbed OS renewal underpinning shortening and OS degeneration in future studies. It is also possible that larger pieces of the OS are being shed and taken up by the RPE. Appreciating why the OSs are breaking down will allow for interventions to halt this process and maintain healthy photoreceptors.

### Killifish as a novel model to explore age-related retinal disease

Killifish are an eminently suitable model to investigate how deposits form at the back of the eye with ageing as well as how these deposits may be impacting retinal and RPE function and health. It is unclear how deposits develop in the ageing human retina and how these deposits impact retinal degeneration. There are few animal models that naturally develop deposits. Zebrafish models with subretinal deposits similar to those observed in ageing killifish here have been reported previously (Noel et al., 2020, 2022; Schlegel et al., 2021). The subretinal deposits observed in killifish appear to be a hybrid deposit; they have features of drusen, as they are lipid rich and oxidized phospholipid containing, but are in the anatomical location of subretinal drusenoid deposits. However, killifish can also develop sub-RPE deposits that are lipid rich, akin to drusen. The association between lipid deposits and OS proteins could be due to the deposits being derived from OS fragments that are not phagocytosed. Alternatively, the accumulating deposits could prevent OS fragments from contacting the RPE, and the OS proteins may be incorporated into deposits after their formation. Future studies can assess whether RPE dysfunction precedes deposit formation or if deposit formation occurs prior to detectable RPE dysfunction. Understanding how and why deposits form will allow for targeted treatments to prevent deposit formation and potentially maintain retinal health for longer periods with age. Fish models of retinal disease have been successfully used for drug screening in the past(Alhasani et al., 2020; Feng et al., 2017; Ganzen et al., 2021; Viringipurampeer et al., 2014; Zhang et al., 2021), and translating these techniques to killifish will provide novel insight into how age-related phenotypes can be influenced.

It is unclear whether killifish display hallmark cellular signs associated with other age-related retinal diseases, such as glaucoma. However, it has been reported that the thickness of the retinal ganglion cell layer significantly declines with age (Vanhunsel et al., 2021). Retinal stretching has been proposed as a means by which the ageing killifish retina thins, rather than neurodegeneration (Bergmans et al., 2023) – however, this study did not investigate the retina for neurodegenerative features, such as opsin mislocalisation. Photoreceptors do not survive post OS loss, and opsin mislocalisation can lead to cell death even when the OS is maintained (Avasthi et al., 2009; Liu et al., 2020; Lopes et al., 2010; Lu et al., 2017; Nakao et al., 2012). Here, the observed signs of degeneration, including OS degradation, infiltration of macrophages, and ceramide accumulation, show clear age-related cellular degeneration in the naturally ageing killifish retina.

### Sex-specific phenotypes in the ageing killifish

We observed sex-specific differences in the ageing killifish retina. The reason for this is unclear but may be due to differences in hormones and metabolism between males and females. Whether there are sex differences in AMD is unclear; some studies suggest that females are more likely to develop late AMD (Owen et al., 2012; Rudnicka et al., 2012), but this may be confounded by females having a longer average lifespan than males, providing more time for AMD to progress in severity. However, a recent study observed that females with intermediate AMD had more complement factor B and I compared to males, suggesting differing molecular responses between the sexes (Marin et al., 2022). Further, a study reported that large drusen and pigmentary changes were more likely to occur in men than women (Sasaki et al., 2018), which coincides with our observation that male killifish had more deposit accumulation than females. It is important to note that detailed histological comparisons between males and females have not been undertaken for AMD, and there may be key distinctions between the sexes not yet identified. Understanding sex-specific differences in disease presentation is valuable for treatment development.

### Future work and conclusions

AMD is a multifactorial condition with a combination of genetics and the environment contributing to an individual’s risk of developing disease. We did not investigate genetic contributions to disease in this study. Our previous transcriptomics study reported that there are transcriptional changes in pathways reported to be linked to AMD, such as oxidative stress and inflammation, as well as changes in AMD linked genes, including *apoeb* and *htra1b* (Bergmans et al., 2024). Subsequent studies will manipulate AMD-related genes in the killifish to introduce human variants and assess how age-related retinal disease is impacted. This would provide important context for why these genetic variants increase risk of developing disease. Further, animals could be exposed to hypoxic environments or fed high fat diets to assess how these factors impact deposit formation and retinal health as the animals get older.

In conclusion, there are many shared pathologies between the human AMD retina and the ageing turquoise killifish retina, including photoreceptor shortening, photoreceptor degeneration, deposit formation, and ceramide accumulation. Additionally, there are sex-specific differences, with males presenting with pathologies earlier and having a more severe phenotype later in life. The killifish provide the opportunity to not only investigate the mechanisms underlying age-related retinal disease, but also the potential to uncover how sex impacts disease.

## Acknowledgements

We thank Moorfields Eye Charity for funding to develop the Institute of Ophthalmology Killifish system (GR001613). This work was supported by a BrightFocus Macular Degeneration Postdoctoral Fellowship (M2022002F), Banting Postdoctoral Fellowship, Moorfields Eye Charity Springboard Award (SB-24B-106), Fight for Sight Seed Grant, Macular Society Seedcorn, and Odette Maymon ECR Research Fund Grant to NCLN. RBM was supported by a BBSRC David Phillips Fellowship (BB/S010386/1) and a BBSRC grant (UKRI705).

## Figures

**Figure S1.**
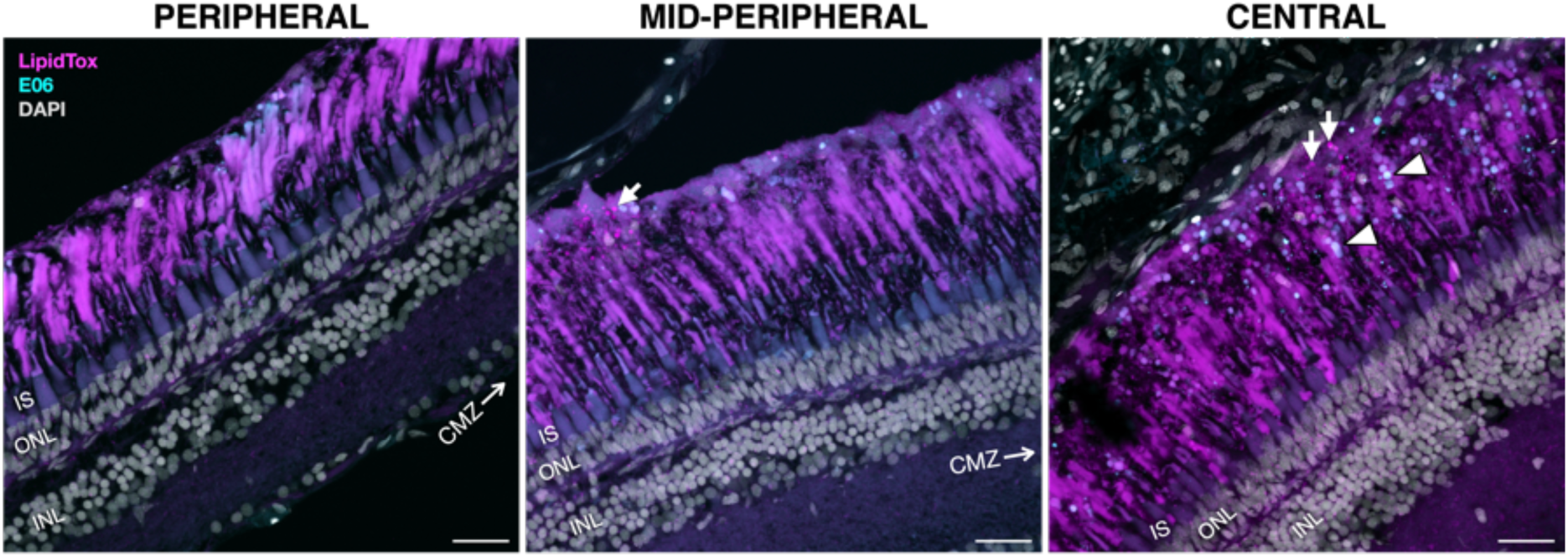
Deposits increase towards the central retina in old killifish. 24-week-old male killifish retina stained with LipidTox (magenta) and oxidised phospholipids (cyan). Peripheral retina has no deposits, while deposits can begin to be observed in the mid-periphery (arrow). In the central retina, an abundance of both neutral lipid (arrows) and oxidised phospholipid-rich deposits (arrowheads) are observed. INL = inner nuclear layer. ONL = outer nuclear layer. CMZ = ciliary marginal zone. Scale bar = 25µm.

**Figure S2.**
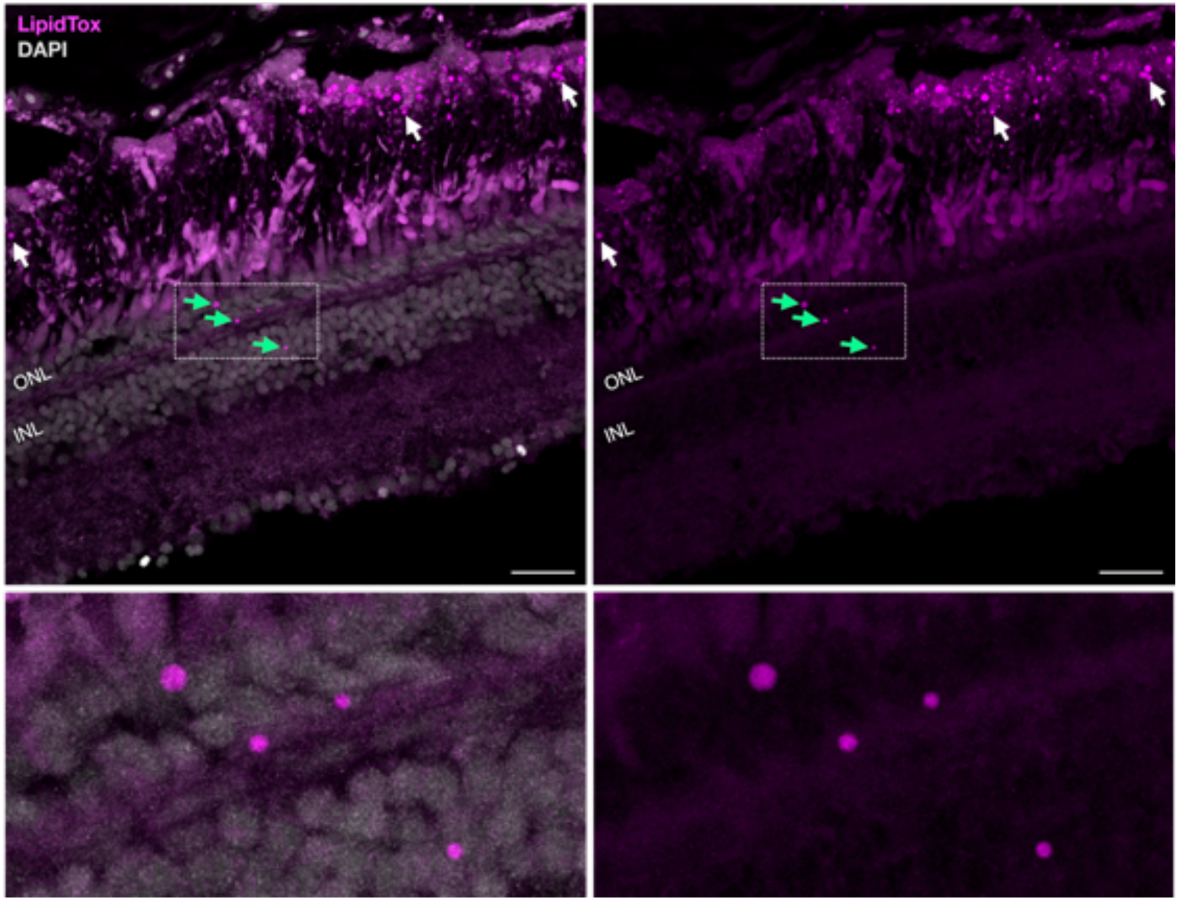
Lipid deposits in the outer and inner nuclear layers of old killifish. Male 24-week-old killifish retina labelled with LipidTox; left panels show merge with DAPI, right panels show only LipidTox. There were lipid deposits between the photoreceptors and the RPE (white arrows) as well as lipid deposits within the ONL and INL (green arrows). Dotted boxes show magnified regions. ONL = outer nuclear layer; INL = inner nuclear layer. Scale bars = 25µm.

**Figure S3.**
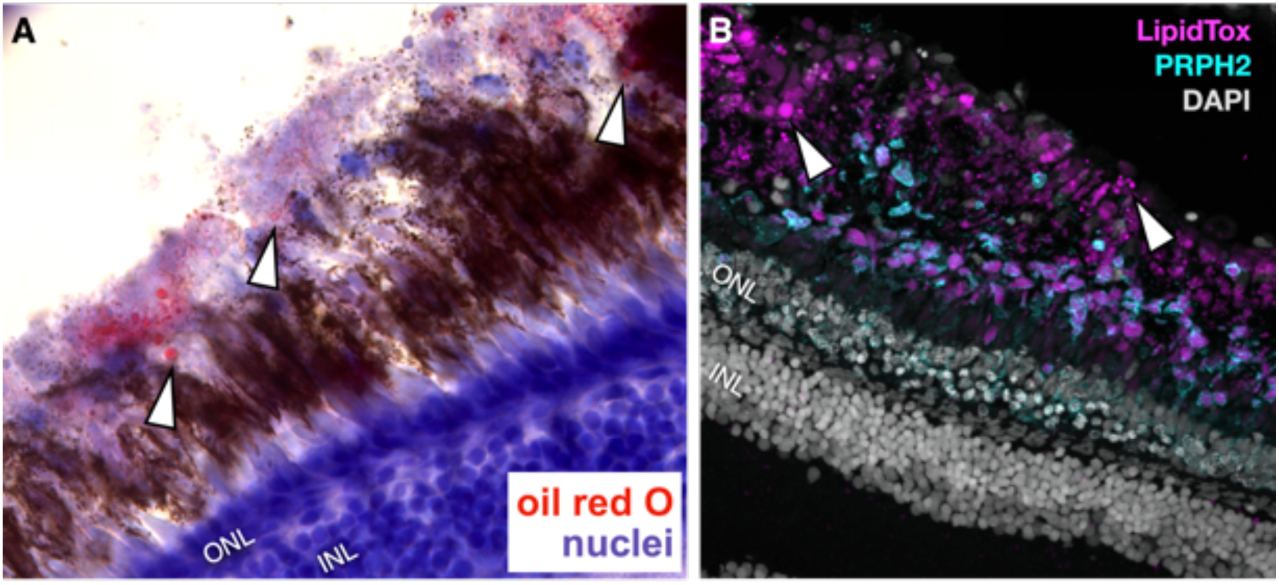
Lipid deposits (arrowheads) can be observed at 18 weeks of age in male killifish. (A) Oil red O (red) and hematoxylin (blue) staining. (B) Neutral lipids (LipidTox, magenta) and outer segment (PRPH2, cyan) labelling. INL = inner nuclear layer. ONL = outer nuclear layer.

**Figure S4.**
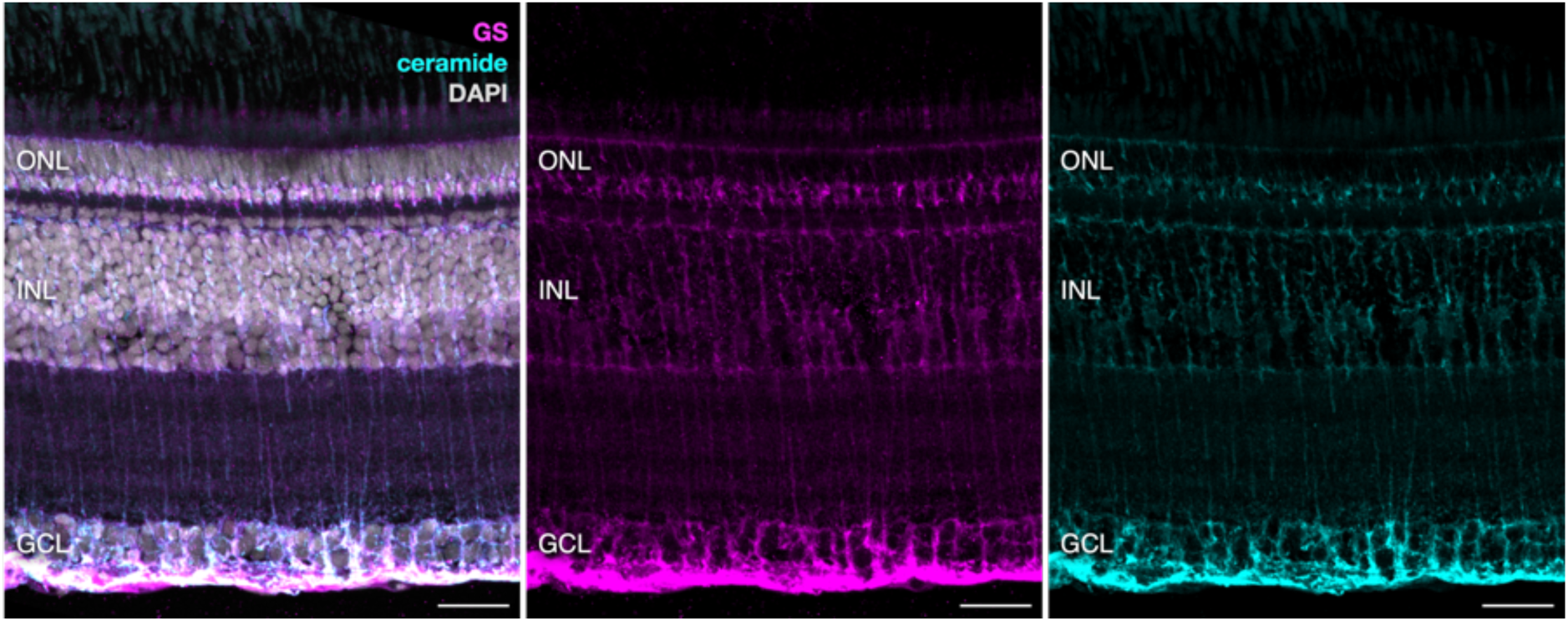
Ceramide localises to killifish Müller glia. 6-week-old killifish retina labelled with glutamine synthetase (GS, magenta) and ceramide (cyan). GCL = ganglion cell layer. INL = inner nuclear layer. ONL = outer nuclear layer. Scale bars = 25µm.

